# Modeling the Evolution of Populations with Multiple Killer Meiotic Drivers

**DOI:** 10.1101/2023.09.28.560003

**Authors:** José Fabricio López Hernández, Boris Y Rubinstein, Robert L. Unckless, Sarah E. Zanders

## Abstract

Meiotic drivers are selfish genetic loci that can be transmitted to more than half of the viable gametes produced by a heterozygote. This biased transmission gives meiotic drivers an evolutionary advantage that can allow them to spread over generations until all members of a population carry the driver. This evolutionary power can also be exploited to modify natural populations using synthetic drivers known as ‘gene drives’. Recently, it has become clear that natural drivers can spread within genomes to birth multicopy gene families. To understand intragenomic spread of drivers, we model the evolution of two distinct meiotic drivers in a population. We employ the *wtf* killer meiotic drivers from *Schizosaccharomyces pombe*, which are multicopy in all sequenced isolates, as models. We find that a duplicate *wtf* driver identical to the parent gene can spread in a population unless, or until, the original driver is fixed. When the duplicate driver diverges to be distinct from the parent gene, we find that both drivers spread to fixation under most conditions. Finally, we show that stronger drivers make weaker drivers go extinct in most, but not all, polymorphic populations with absolutely linked drivers. These results reveal the strong potential for natural meiotic drive loci to duplicate and diverge within genomes. Our findings also highlight duplication potential as a factor to consider in the design of synthetic gene drives.

## INTRODUCTION

Most alleles are Mendelian in that they are transmitted to half of the offspring of a given individual. Meiotic drive alleles, in contrast, can be passed on to more than half, even all offspring. Meiotic drive is a powerful evolutionary force as the transmission bias allows a meiotic driver to spread in a population (Novitski 1957). Understanding the spread of meiotic drivers within populations is critical for deciphering the evolution of natural populations and may guide design of synthetic gene drives that aim to control natural populations (Price et al. 2020; Lindholm et al. 2016 ; Zanders and Unckless. 2019).

The evolution of single drive loci in populations has been extensively modeled (Bull 2016; Crow 1991; Dyer and Hall 2019; Fishman and Kelly 2015; Hall and Dawe 2018; Hartl 1970; Lopez Hernandez et al. 2021; Manser et al. 2020; Martinossi-Allibert et al. 2021). However, some species carry multiple, unrelated meiotic drivers (Akera et al. 2019; Bravo Nunez et al. 2020a; Cazemajor et al. 2000; Dalstra et al. 2003; Didion et al. 2015; Long et al. 2008; Lyon 2003; Tao et al. 2007a; Voelker and Kojima 19971; Vogan et al. 2019; Yang et al. 2012, Yu et al. 2018). Additionally, some drive genes are members of multi-gene families (Dawe et al. 2018; Hu et al. 2017; Muirhead and Presgraves 2021; Nuckolls et al. 2017; Vedanayagam et al. 2021; Vogan et al 2019). One potential evolutionary implication of species carrying multiple distinct allelic drivers has recently been explored using evolutionary modeling (Bravo Nunez et al. 2020b). However, the evolution of populations polymorphic for multiple drivers born from gene duplication has not been formally considered.

The *wtf* killer meiotic drivers found in fission yeasts (*Schizosaccharomycetes*) have undergone many gene duplication events over the past ∼119 million years (Figure 1A; De Carvalho et al. 2022). In *S. pombe*, distinct isolates encode between 4-14 genes that appear to be intact drivers (Eickbush et al. 2019; Hu et al. 2017). Each *wtf* driver encodes a poison and an antidote protein from separate, but largely overlapping transcripts of the same gene. All four developing meiotic products (spores) are exposed to the poison, while only those that inherit the driving *wtf* gene acquire enough antidote to neutralize the poison (Figure 1B; Hu et al. 2017; Nuckolls et al. 2017, Nuckolls et al. 2022). Importantly, the antidotes encoded by a given *wtf* driver generally provide no protection against the poisons of distinct drivers with different sequences (Bravo Nunez et al. 2020b; Hu et al. 2017).

**Figure 1.**
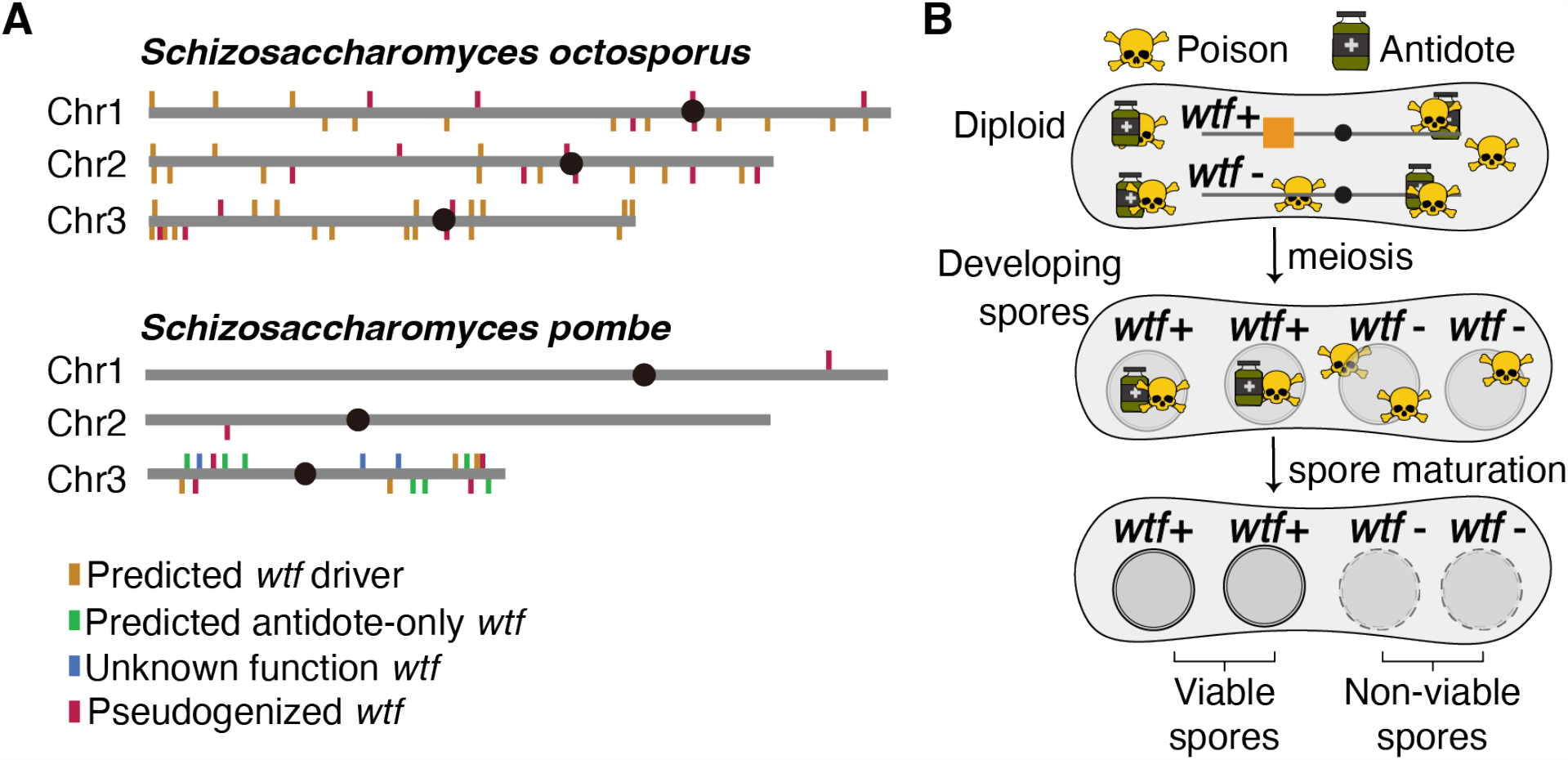
Poison-antidote *wtf* meiotic drivers in *Schizosaccharomyces*. **A**) Genomic loci that contain members of the *wtf* gene family in *S. octosporus* and *S. pombe* reference genomes (Eickbush et al. 2019; De Carvalho 2022). Each marked locus contains at least one of the indicated *wtf* genes. **B)** In *S. pombe*, a *wtf* meiotic driver produces both a poison and an antidote that are expressed in diploids induced to undergo meiosis. After meiosis, the antidote is enriched only in the spores that inherit the driver. The antidote rescues only the cells that inherit the driver, while the rest of the gametes are susceptible to the poison.

Here, we model the evolution of meiotic driver gene duplicates using parameters derived from the *wtf* drivers. We find that duplicates of *wtf* drivers are likely to spread in a population under many conditions, particularly when one of the duplicate genes diverges to become a distinct driver. Our results help explain the evolution of an ancestral *wtf* driver into a gene family and suggest that compact poison-antidote drivers are prone to spreading within genomes.

## METHODS

### Model for identical *wtf* drivers

When starved, haploid *S. pombe* cells can mate to form a diploid that undergoes meiosis to produce four haploid progeny, known as spores (Forsburg and Rhind 2006). While the relative time spent in the haploid phase is different from diploid eukaryotes, the same types of equations can be used to model allele frequency changes over generations of sexual reproduction (Crow 1991; Lopez Hernandez et al. 2021).

We initially modeled the evolution of a pair of identical *wtf* driver duplicates, *wtfB* and *wtfA*, at distinct, but linked loci over successive rounds of sexual reproduction. Our equations are extensions of those presented in Crow 1991. Each driver has only one alternate allele that does not drive (e.g., *wtfA*-). A total of four distinct genotypes are therefore possible: *wtfA*+ *wtfB*+, *wtfA*+ *wtfB*-, *wtfA*- *wtfB*+, and *wtfA*- *wtfB*-. Those genotypes are found with frequencies *x*_1_, *x*_*2*_, *x*_*3*_, and *x*_*4*_, respectively (Table 1). We assume an infinitely large population, equal fitness of all haploid genotypes during clonal growth, and random mating.

**Table 1.** Parameters and variables used in the modeling of two drivers in a fission yeast population.

| Parameters/variables | Description | Parameter range |
| --- | --- | --- |
| $x_1$ | Frequency of genotype $wtfA+$ $wtfB+$ | 0 to 1 |
| $x_2$ | Frequency of genotype $wtfA+$ $wtfB-$ | 0 to 1 |
| $x_3$ | Frequency of genotype $wtfA-$ $wtfB+$ | 0 to 1 |
| $x_4$ | Frequency of genotype $wtfA-$ $wtfB-$ | 0 to 1 |
| $t$ | Transmission advantage | 0 to 1 |
| $r$ | Recombination frequency between $wtf$ loci | 0 to 0.5 |

As the drivers are identical, drive (spore killing) will only affect spores that inherit no *wtf*+ alleles from a diploid carrying one or more *wtf*+ alleles (Figure 2A). The parameter ‘*t*’ is the fraction of spores killed by a driver, so it thus represents the strength of drive (Table 1). Spores that inherit neither *wtfA*+ or *wtfB*+ from a diploid heterozygous for both are susceptible to killing by both drivers (i.e., a fraction represented by *2t* − *t*^*2*^ are killed (and (1 − *t*)^*2*^ survive) since *t*_*A*_ *= t*_*b*_ *= t*). We assign no additional fitness costs to any genotypes, beyond the costs caused by the driver due to spore killing.

**Figure 2.**
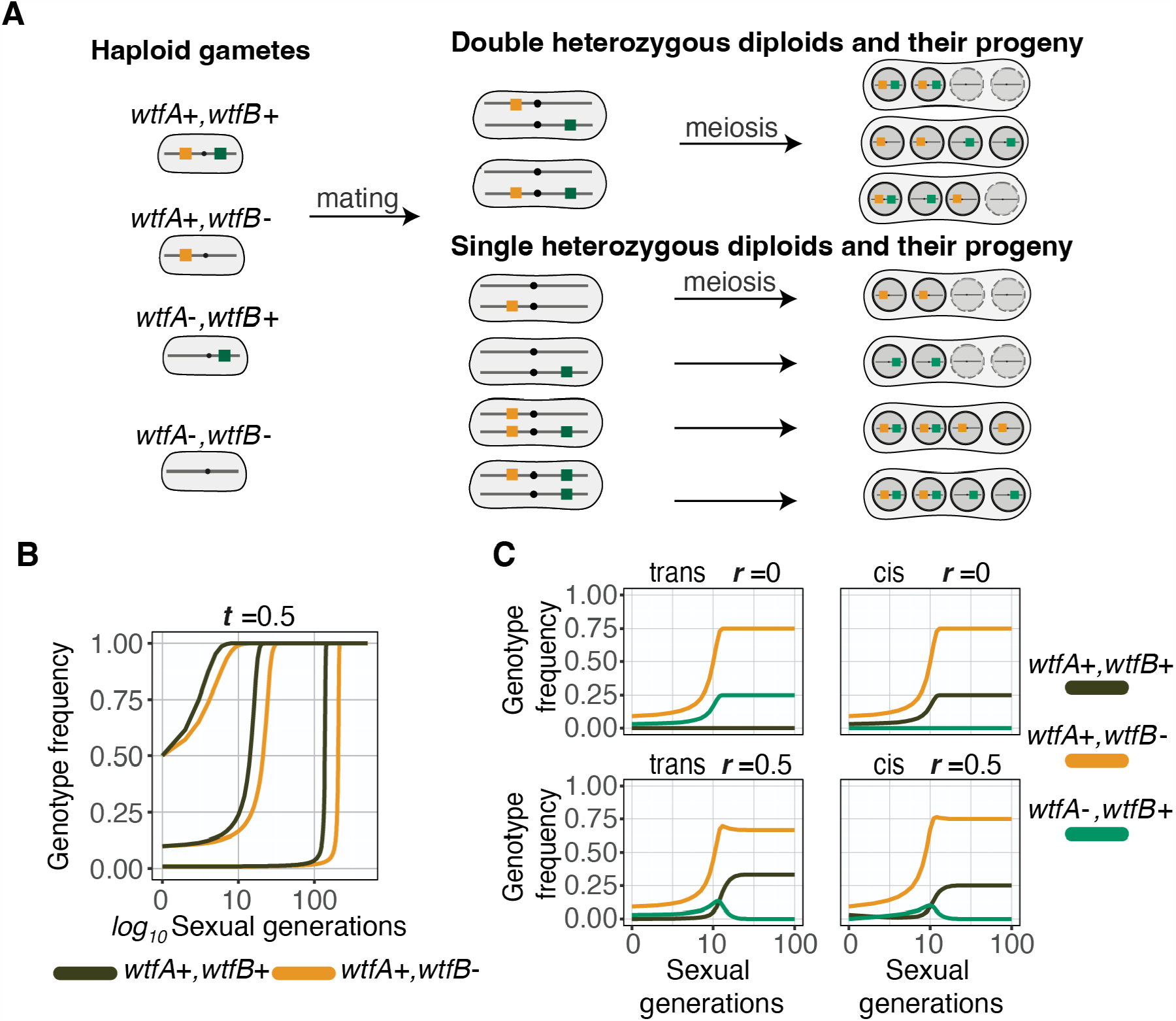
Evolution of two identical drivers after gene duplication. **A**) Four distinct genotypes are possible in the population after a *wtf* driver (*wtfA+*) duplicates: *wtfA*+ *wtfB*+, *wtfA*+ *wtfB*-, *wtfA*- *wtfB*+ and *wtfA*- *wtfB*-. Those haploids can mate to form diploids with a variety of genotypes. Drive will occur if diploids are heterozygous for one or two drivers. Spores that do not inherit one of the drivers from a heterozygote are susceptible to killing. Live spores are shown within a solid black circle whereas spores susceptible to killing by drive are shown within a dotted circle. **B)** Simulations between two populations with only one driver (*wtfA*+ *wtfB*-, orange) or two drivers absolutely linked in cis (*wtfA*+ *wtfB*+, black) spreading in a popullaion where the alternate genotype lacks drivers (*wtfA*- *wtfB*-). The initial frequencies of the *wtfA*+ *wtfB*- and *wtfA*+ *wtfB*+ genotypes shown are 0.01, 0.1, and 0.5. The transmission advantage (*t*) for each driver is 0.5. **C)** Four distinct simulations in which a driver (*wtfA*+) makes an identical duplicate (*wtfB+*) in trans (on the homologous chromosome, left) or in cis (on the same chromosome, right). The transmission advantage (*t*) for each driver is 1. Simulations where the duplicate gene is absolutely linked (*r*=0, top) and unlinked (*r*=0.5, bottom) from the parent gene are shown. The starting frequency of the ancestral genotype (*wtfA*+ *wtfB*-, orange) is 0.1. The starting frequency of genotypes with a duplicated driver in cis (*wtfA*+ *wtfB*+, black) or in trans (*wtfA*- *wtfB*+, green) is 0.03. The remainder of each population is comprised of the *wtfA*- *wtfB*- genotype.

The frequency at which recombinant genotypes form in the spore of double heterozygotes (e.g. *wtfA*+ *wtfB*+/*wtfA*- *wtf-)* diploids is determined by ‘*r*’ (Table 1). To simplify calculating genotype frequency changes due to recombination during gametogenesis, we use the parameter ‘*D*’ where *D = r*(*x*_1_*x*_*4*_ − *x*_*2*_*x*_*3*_) (Crow, 1991). The frequency of each genotype in subsequent generations of a given starting population can be calculated using the equations 1.1 through 1.4. Each equation includes a parameter for population fitness 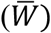, which is defined in equation 1.5. To calculate the frequency of *wtfA*+ *wtfB*+ spores in the next generation (*x′*_1_), we considered all possible diploid genotypes that can generate *wtfA*+ *wtfB*+ spores:

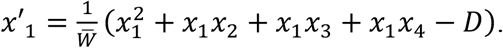

The equation can be rewritten as:

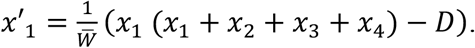

After considering that *x*_1_ + *x*_*2*_ + *x*_*3*_ + *x*_*4*_ *=* 1, we can further simplify to

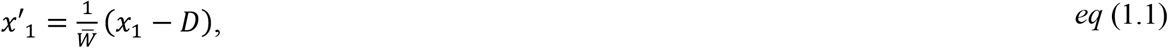

To calculate the frequency of the *wtfA*+ *wtfB*- and *wtfA*- *wtfB*+ genotypes in the next generation (*x′*_*2*_ and *x′*_*3*_, respectively), we can use very similar equations as that used to calculate *x′*_1_, however, we use ‘+*D*’ to reflect the change in the two genotypes due to recombination as follows:

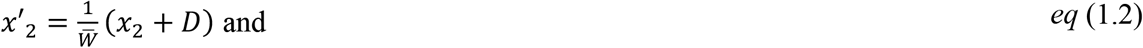

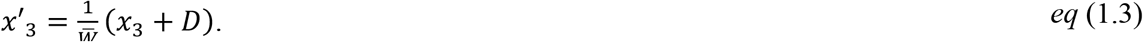

To calculate the frequency of spores with the *wtfA*- *wtfB*- genotype in the next generation (*x′*_*4*_), we must consider that those spores are susceptible to killing. When *wtfA*- *wtfB*- spores are generated by a single heterozygote, (1 − *t*) spores survive, whereas (1 − *t*)^*2*^ *wtfA*- *wtfB*- spores survive when they are generated by a diploid heterozygous for both drivers. Considering the fitness of each diploid, we can calculate *x*’_4_ with the equation below.

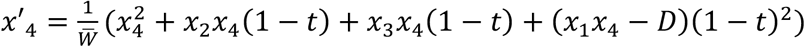

rewritten as,

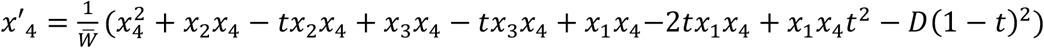

or

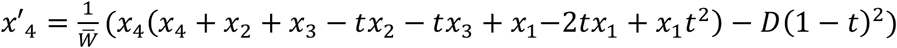

and after considering that *x*_1_ + *x*_*2*_ + *x*_*3*_ + *x*_*4*_ *=* 1, the equation can be simplified to:

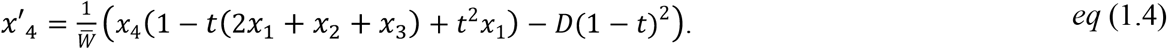

To calculate the mean population fitness 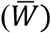, we used the surviving spores produced by all genotypes and their frequencies. The fitness of the *x*_1_, *x*_*2*_ and *x*_*3*_ genotypes is 1. For *x*_*4*_, a fraction of spores are destroyed by drive. The derivation of the fitness of *x*_4_ spores is taken from equation 1.4 described above.

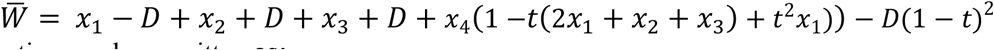

This equation can be rewritten as:

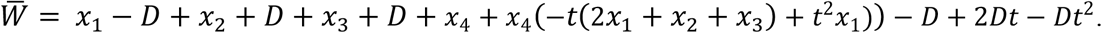

Given that *x*_1_ + *x*_*2*_ + *x*_*3*_ + *x*_*4*_ *=* 1, the equation can be simplified to:

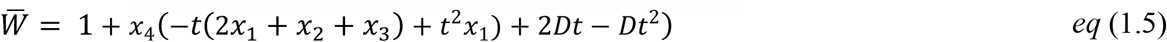

### Model for distinct *wtf* drivers

The model for two distinct *wtf* drivers (again represented by *wtfA*+ and *wtfB*+) uses the same parameters and is similar to the model for identical *wtf* drivers (described above) with one important difference. Namely, spores that inherit *wtfA+* are *not* protected from killing by *wtfB+* and vice versa. Thus, if a diploid is heterozygous for both drivers, a spore must inherit both to be resistant to killing. Because of this, the fitness components of the equations to calculate *x′*_*2*_ through *x′*_*4*_ change as described below.

To calculate the frequency of *wtfA*+ *wtfB*- spores in the next generation (*x′*_*2*_), we calculated that the *wtfB*+ driver will kill a fraction of *wtfA*+ *wtfB-* spores (described by *t*_*B*_) generated by diploids heterozygous for *wtfB* as shown below.

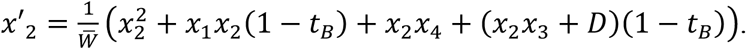

This equation can be rewritten as

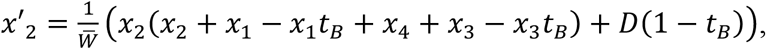

further simplified to:

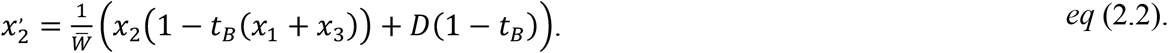

The equation for calculating the frequency of *wtfA*- *wtfB*+ spores in the next generation (*x′*_*3*_), must be similarly amended to include that the *wtfA*+ driver will kill a fraction of *wtfA*- *wtfB+* spores (described by *t*_*A*_) generated by diploids heterozygous for *wtfA* as shown below.

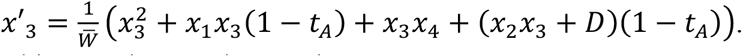

This equation can be rewritten as

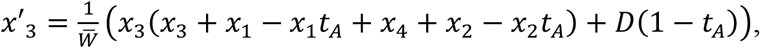

and further simplified to:

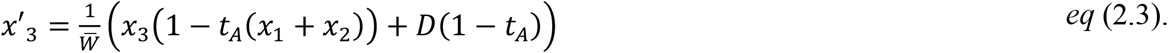

To calculate the frequency of *wtfA*- *wtfB*- spores in the next generation (*x′*_*4*_), we modified the equation to reflect that these spores are sensitive to being killed by *wtfA+* in *wtfA*+ *wtfB*-/*wtfA*- *wtfB*- diploids, by *wtfB*+ in *wtfA*- *wtfB*-/*wtfA*- *wtfB*+ diploids, and by both drivers in diploids heterozygous for both drivers as shown below.

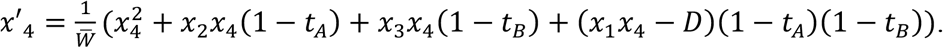

This equation can be expanded to

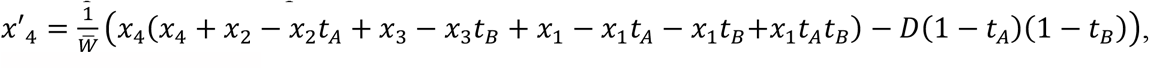

and simplified to

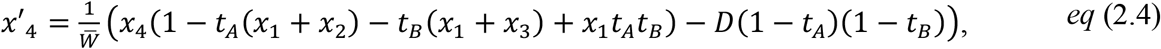

The mean population fitness is again calculated by considering the fitness of all genotypes in the population as follows:

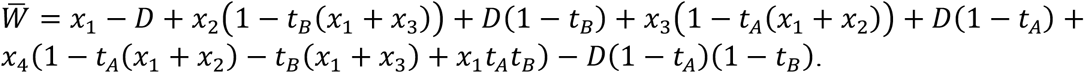

This equation can be expanded to

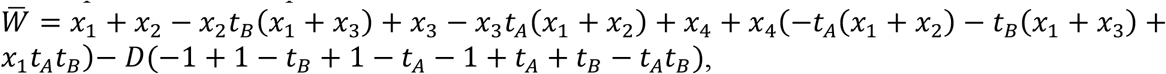

simplified to

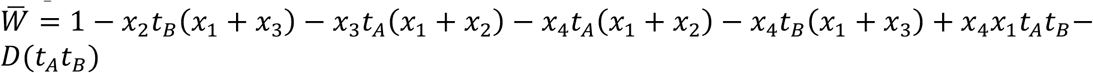

and further simplified to

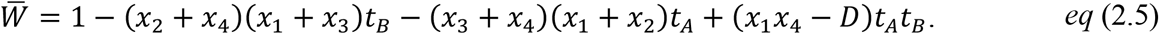

### Model for four distinct drivers on competing haplotypes

To model the evolution of two to four distinct driver alleles at a single locus, we assumed no recombination between drivers (*r =* 0). We designated four possible driver alleles: *wtfA*^*1*^, *wtfA*^*2*^, *wtfA*^*3*^, and *wtfA*^*4*^, with the relative frequencies 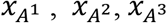 and 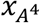, respectively. The spore killing caused by each driver is defined by the *t* value for that driver. Drive will occur in heterozygotes such that each spore is susceptible to being killed by the driver it does not inherit. Drive does not, however, occur in homozygotes.

The frequency of each allele in subsequent generations can be calculated as follows:

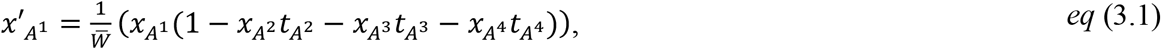

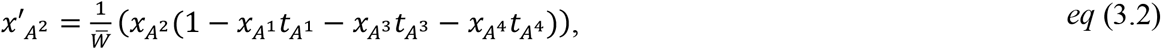

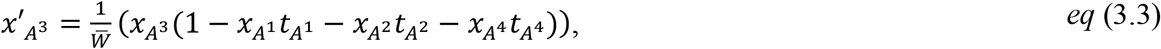

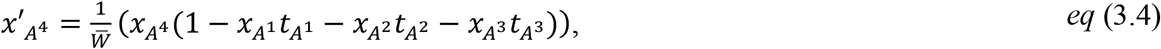

where mean population fitness was

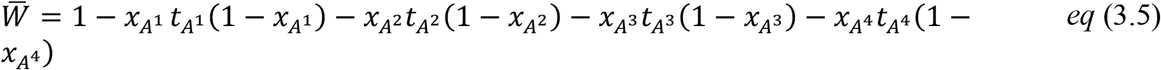

### Simulations, stability analysis, data analysis, and visualization

We carried out deterministic forward simulations using a range of starting genotype frequencies. For simulations where drivers enjoyed the same transmission advantage, we explored the full parameter space of distinct initial genotype frequencies, and we varied *t* from 0 to 1 with a 0.05 step size. For simulations where the two drivers had distinct transmission advantages, we first randomly sampled 35,370 from the 176,676 possible initial genotype frequencies that range from 0 to 1 with a 0.01 step size. Using the sampled frequencies, we assigned 5000 distinct combinations of transmission advantages ranging from 0.01 to 1, with 0.01 step to each set of initial genotype frequencies considered. This approach generated a total of 176.775 million possible initial conditions. From those conditions, we randomly sampled 10,000 and simulated them.

For all simulations, we assumed an infinitely large population and simulated 10,000 generations or until a steady state with a genotype frequency change less than 1×10^−15^ occurred. For each generation, we tracked and updated the genotype frequencies and mean population fitness. To determine the fate of each genotype, all frequencies close to 1 or 0 were rounded with tolerance of 1×10^−13^, lower than the inverse reported effective population size 1×10^5^ -1×10^9^ (Farlow et al. 2015; Tusso et al. 2019). A genotype was considered fixed when it equaled 1 and extinct when it equaled 0.

We also determined possible genotype frequency steady state solutions using Mathematica (Wolfram Research 2021). We defined the steady state of the recurrence equations by identifying that the equations follow the form, 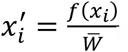. Where 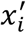 is the frequency of each genotype “*i*” to the next generation which depends on the mean population fitness 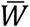 and a function of the absolute frequency of each genotype *f*(*x*_*i*_). The steady state is determined by the condition in which the change of all genotypes to the next generation equals 0, 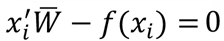.

Steady state solutions were be determined by simplifying the system of equations to *x*_*4*_ *=* 1 − *x*_1_ + *x*_*2*_ + *x*_*3*_. Solutions were found for the cases *r =* 0 or *t*_*A*_ *= t*_*B*_ including a particular case were *t*_*A*_ *= t*_*B*_ *=* 1.

To determine the mathematical stability of the solutions, we used the eigenvalues of the Jacobian matrix for all four recurrence equations (Otto and Day. 2007). A solution is stable only when the leading eigenvalue is less than 1 and unstable when it is greater than 1. In cases where the associated eigenvalues are exactly one, the solution stability cannot be defined by the Jacobian matrix alone. The solution when the *wtfA*+ *wtfB*+ genotype is fixed not defined by the Jacobian except upon perturbation of the genotype frequencies (Table 2, see Suppl. material for mathematical proof).

**Table 2.**
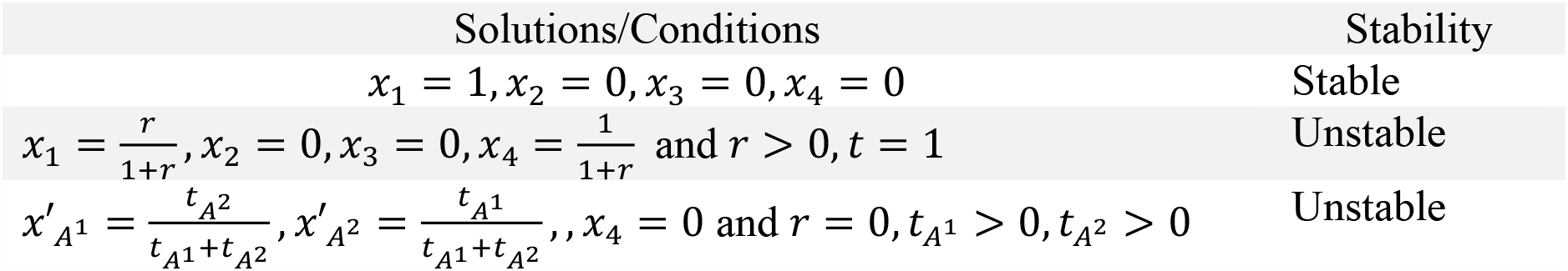
The solutions and stability associated to leading eigenvalues for two distinct drivers.

| Solutions/Conditions | Stability |
| --- | --- |
| $x_1 = 1, x_2 = 0, x_3 = 0, x_4 = 0$ | Stable |
| $x_1 = \frac{r}{1+r}, x_2 = 0, x_3 = 0, x_4 = \frac{1}{1+r}$ and $r > 0, t = 1$ | Unstable |
| $x'_{A^1} = \frac{t_{A^2}}{t_{A^1}+t_{A^2}}, x'_{A^2} = \frac{t_{A^1}}{t_{A^1}+t_{A^2}}, x_4 = 0$ and $r = 0, t_{A^1} > 0, t_{A^2} > 0$ | Unstable |

All simulations were coded in R (Team 2019. Version 4.2.3) and performed using R. The code is available at: https://github.com/ZandersLab/Genetic_conflict_between_poison-antidote_meiotic_drivers

## RESULTS

### Evolution of two identical *wtf* paralogs

Initially after a gene duplication, the two paralogs are likely to be identical. We thus first considered the evolution of two identical drivers: *wtfA* and its paralog, *wtfB*. Briefly, our model considers an infinitely large population, random mating, and no fitness costs beyond the fraction of spores destroyed by drive. There are four haploid genotypes possible: *wtfA*+ *wtfB*+, *wtfA*+ *wtfB*-, *wtfA*- *wtfB*+, and *wtfA*- *wtfB*-. Because *wtfA* and *wtfB* are identical, drive will occur in diploids that are: 1) heterozygous for both drivers and 2) in diploids heterozygous for one driver and lacking the second driver. In both cases, only spores that that do not inherit either driver (*wtfA- wtfB-)* can be destroyed by drive (Figure 2A). We use the term ‘*t*’ to reflect the transmission advantage of each driver in heterozygotes. For example, at *t =* 1, all *wtfA- wtfB-* spores produced by diploids heterozygous for both drivers would be destroyed. At, *t =* 0.5, 75% (*2t* − *t*^*2*^) of the *wtfA- wtfB-* spores from diploids heterozygous for both drivers are destroyed. We used the parameter ‘*r*’ to reflect recombination frequencies with *r =* 0 and *r =* 0.5 for absolutely linked and unlinked loci, respectively.

We modeled populations with varying starting frequencies of the four haploid genotypes. We found that two identical drivers with 0 < *t* < 1 that are absolutely linked in cis (*wtfA*+ *wtfB*+) spread in a population of *wtfA*- *wtfB*- cells faster than a haplotype containing a single driver locus (*wtfA*+ *wtfB*-) (Figure 2B). As described above, if *wtfA* has a transmission advantage of *t =* 0.5, and generates a tandem duplicate *wtfB*, that effectively generates a drive locus equivalent to a single driver with *t =* 0.75. If a driver with the strongest possible transmission advantage (*t =* 1) makes a tandem duplicate, the dynamics of driver spread are the same as if the duplicate did not occur.

Overall, while the *wtfA*- *wtfB*- genotype is present in a population, drive can occur and both drive alleles (*wtfA+* and *wtfB+*) spread. After fixation of *wtfA*+, drive no longer occurs and allele frequencies remain constant. The frequency of each haplotype when fixation of the original driver (*wtfA*+) occurs can vary depending on recombination and whether the *wtfB*+ duplicate occurs in cis (to generate *wtfA*+ *wtfB*+) or in trans to generate *wtfA*- *wtfB*+ (Figure 2C).

### Evolution of two distinct *wtf* paralogs

We next considered the evolution of a pair of *wtf* paralogs (*wtfA+* and *wtfB+*) when one of the pair diverges such that they become distinct, mutually-killing drivers. Because the drivers are distinct, drive will occur in diploids heterozygous for one or both drivers (Figure 3A). The *wtfA+* and *wtfB+* drivers will destroy a fraction of spores that do not inherit them from heterozygotes determined by the parameters ‘*t*_*A*_*’* and ‘*t*_*B*_*’*, respectively. When a single driver is heterozygous, the fraction of dead spores is determined by only one *t* parameter. When both drivers are heterozygous, the fraction of spores not inheriting both drivers that survive will be determined by both *t*_*A*_ and *t*_*B*_ (Figure 3B).

**Figure 3.**
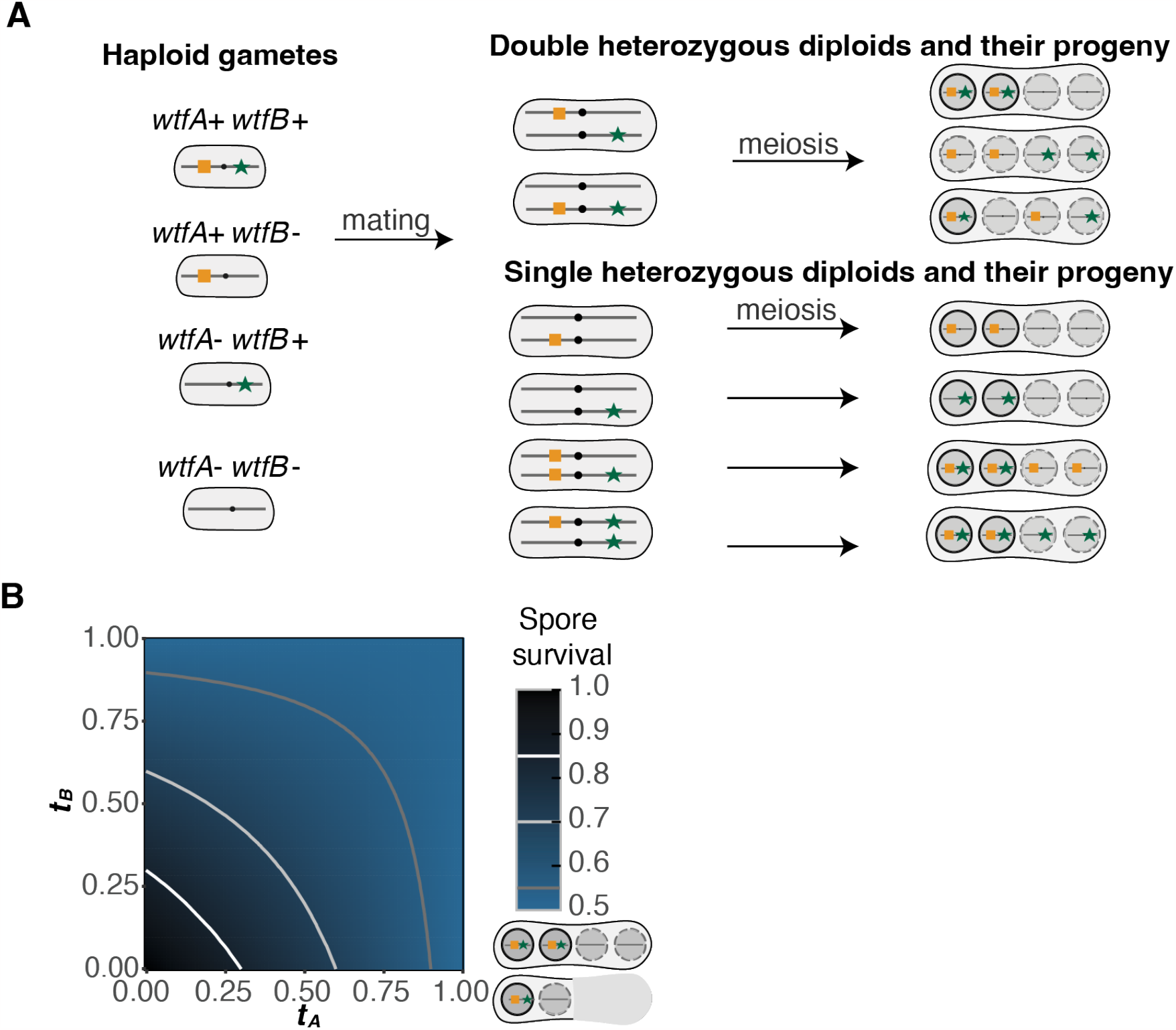
Spore survival with two distinct *wtf* meiotic drivers in a population. **A**) Cartoon of the four possible genotypes that carry one (*wtfA*+ *wtfB-* and *wtfA- wtfB*+), two (*wtfA*+ *wtfB*+) or no (*wtfA*- *wtfB-*) meiotic drivers. Haploids can mate to form diploids of a variety of genotypes, including heterozygotes which are illustrated. Drive will occur in the diploids shown as spores are susceptible to being killed by each driver they do not inherit from a heterozygote. Live spores are shown within a solid black circle whereas spores susceptible to killing by drive are shown within a dotted circle. **B)** The fraction of spores produced by diploids heterozygous for two drivers that inherit two or zero drivers expected to survive when considering varying drive strength.

We first considered populations with the two drivers on the same chromosome (linked in cis or trans), and in the absence of recombination (*r*=0). We also initially assumed that the two drivers were of equal strength (*t*_*A*_ *= t*_*B*_ *= t*). We considered populations with varying starting frequencies of each genotype ranging from 0 to 0.95. We found that in all populations that initially included individuals of the *wtfA*+ *wtfB*+ genotype, that genotype eventually drove to fixation (Figure 4A). Consistently, the fixation of that double driver genotype occurred faster when the drivers were stronger (Figure 4A). The magnitude of the changes in the *wtfA*+ *wtfB*+ genotype frequencies between generations varies depending on frequencies of the remaining genotypes. This is easy to visualize when one considers a population with only three genotypes (*wtfA*+ *wtfB*+, *wtfA*+ *wtfB*- and *wtfA*- *wtfB*-), as would occur if the duplication event creating *wtfB*+ happened once on the same haplotype as *wtfA*+ (Figure 4B). In this scenario we see that the drive alleles have a larger frequency change between generations, when many driver heterozygotes are likely to be formed by mating (Figure 4B). The fixation of the genotype *wtfA*+ *wtfB*+ is ensured once it is present in the population (Table 2; see proof in the Suppl. material).

**Figure 4.**
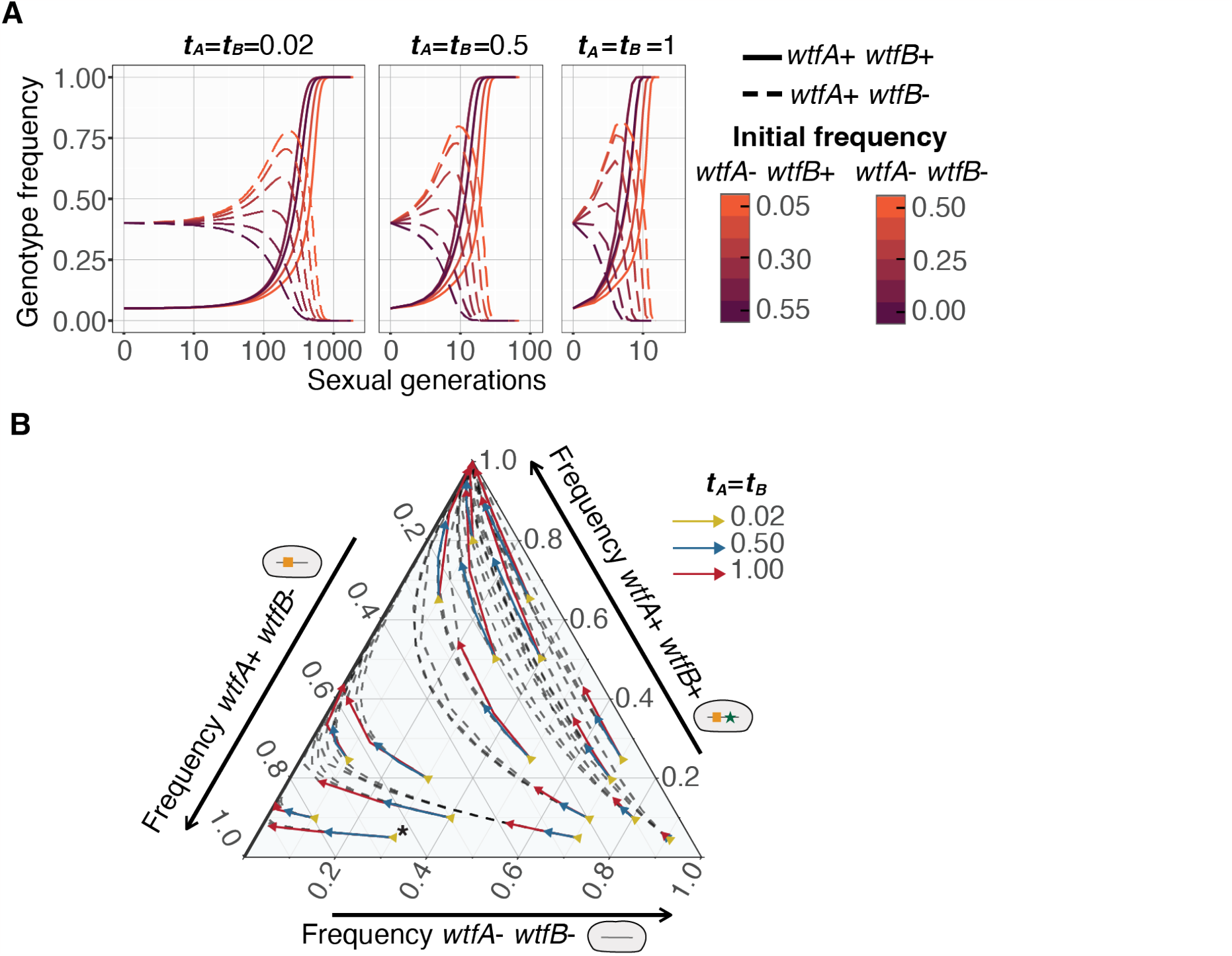
The evolution of populations with two drivers of equal strength in the absence of recombination. **A**) Change in driver genotype frequencies over time. The genotype frequencies of *wtfA+ wtfB*+ (solid, 0.05 initial frequency), *wtfA+ wtfB-*(dashed, 0.40 initial frequency) with varying *wtfA- wtfB-* initial frequencies with 0.1 steps. The remainder of each population is comprised of *wtfA*- *wtfB*+ genotype. The genotype *wtfA+ wtfB+* goes to fixation when present. Strong drivers (*t =* 1, right) spread to fixation faster than weak drivers (*t =* 0.0*2*, left). **B)** The evolution of populations lacking the *wtfA+ wtfB-* genotype but with varying frequencies of the other genotypes is shown. To read the frequency of the *wtfA+ wtfB+* genotype, follow a horizontal line to the right axis. To read the frequency of the *wtfA- wtfB-* genotype, follow the diagonal down and to the left to the bottom axis. To read the frequency of the *wtfA+ wtfB-* genotype, follow the diagonal up and to the left to the left axis. Every point in the triangle represents the frequency in three axes that add up to 1. The point marked with an * represents the following frequencies: *wtfA*- *wtfB*- of 0.30, *wtfA+ wtfB*+ of 0.05, and *wtfA*+ *wtfB*- of 0.65. The arrows depict allele frequency changes over four generations from that starting point and the dotted lines show subsequent frequency changes. Arrow color reflects drive strength (*t*) as shown in the key and overlapping arrows share a common starting point. In the absence of one single driver genotype (*wtfA- wtfB+*), the genotype with both drivers (*wtfA+ wtfB+*) spreads to fixation. When the initial frequency of the double driver genotype (*wtfA+ wtfB+*) is low, the single driver genotype (*wtfA+ wtfB-*) can increase in frequency for a few generations (lines moving left), due to outcompeting the driverless genotype (*wtfA- wtfB-)*, prior to being replaced by the double driver genotype (arrows pointing up).

We next considered the evolution of drivers of equal strength (*t*_*A*_ *= t*_*B*_) in the presence of recombination, *r* > 0. We found that in almost all cases, both drivers spread to fixation. As before, stronger drivers reach fixation faster (Figure 5A-5E). Interestingly, in some cases the frequency of the double driver genotype (*wtfA*+ *wtfB*+) initially decreases prior to increasing to spread to fixation (Figure 5B, 5E). This occurs when the frequency of the *wtfA*- *wtfB-* is relatively high and the *wtfA*+ *wtfB*+ frequency is relatively low, following the condition 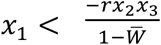. In such cases, double heterozygotes are created and progeny can form recombinant spores that inherit a single driver are thus destroyed by the opposite driver. Strikingly this effect can even lead to loss of the double driver genotype when drive is strong (*t =* 1), no single driver genotypes is present, and the double driver genotype has a low initial frequency (Figure 5F; Table 2).

**Figure 5.**
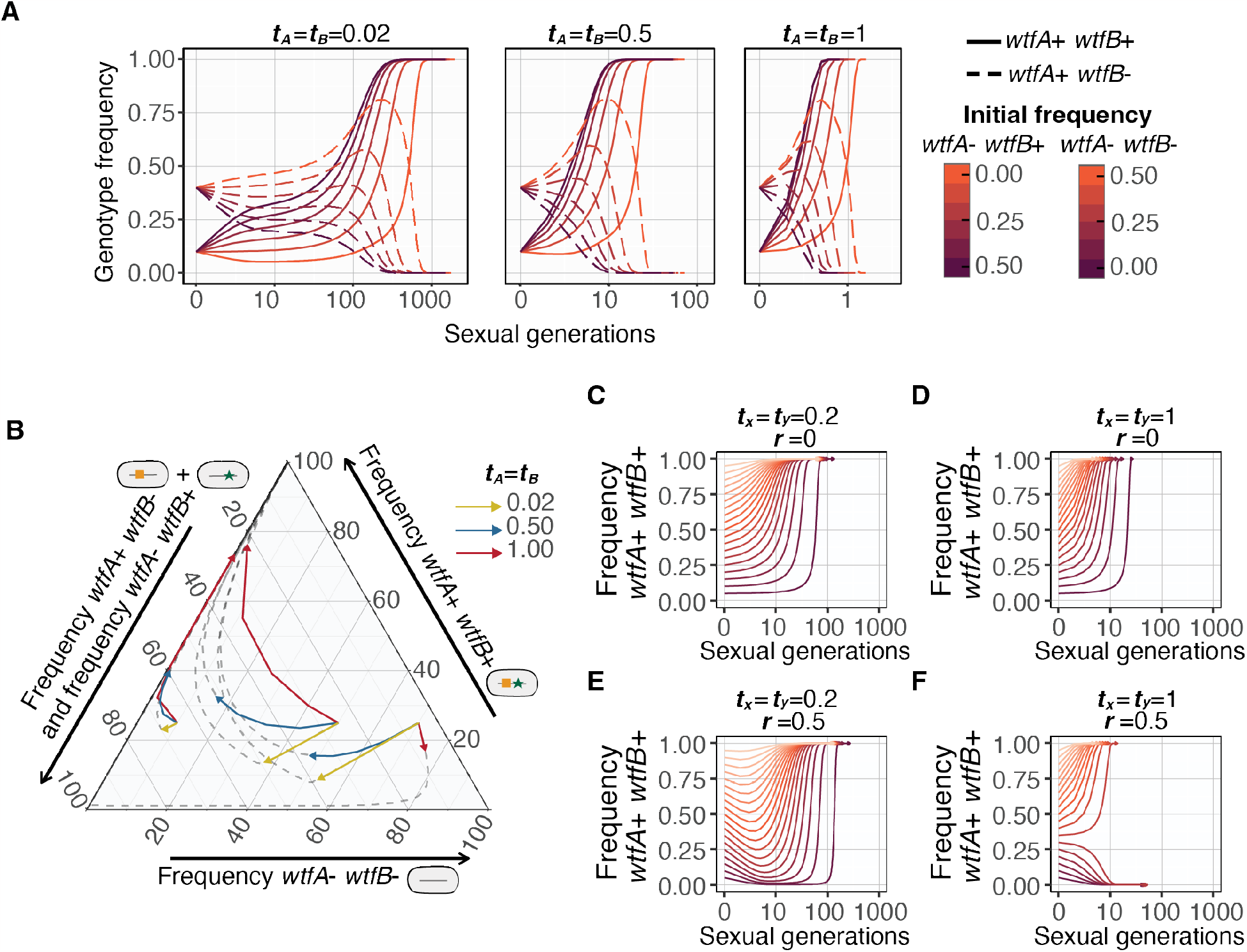
The evolution of populations with two drivers of equal strength in the presence of recombination. **A**) Changes in two unliked driver genotypes over time (*r =* 0.5). The genotype frequencies of *wtfA+ wtfB*- (solid, 0.1 initial frequency), *wtfA+ wtfB+* (dashed, 0.40 initial frequency) with varying *wtfA- wtfB-* initial frequencies with 0.1 steps. The remainder in each population is comprised of genotype *wtfA*- *wtfB*+. Strong drivers (*t =* 1, right) spread to fixation faster than weak drivers (*t =* 0.0*2*, left). **B)** The evolution of populations initially lacking the *wtfA*- *wtfB*+ genotype. The frequency of each genotype is shown on the three axes. To read the frequency of the *wtfA+ wtfB+* genotype, follow a horizontal line to the right axis. To read the frequency of the *wtfA- wtfB-* genotype, follow the diagonal down and to the left to the bottom axis. To read the combined frequency of the *wtfA+ wtfB-* and *wtfA- wtfB+* genotypes, follow the diagonal up and to the left to the left axis. The two unlinked drivers have equal strength and three driver strengths (indicated by the different arrow colors as shown in the key) were considered. The arrows depict allele frequency changes over four generations from that starting point and the dotted lines show subsequent frequency changes. Although the frequency of the *wtfA*+ *wtfB*+ genotype can initially decline (downward arrows), that genotype eventually spreads to fixation under all conditions illustrated. **C-F)** Four simulated populations initially carry only two genotypes (*wtfA*+ *wtfB*+) and (*wtfA*- *wtfB*-). The initial frequencies for the genotype *wtfA*+ *wtfB*+ range from 0.05 to 0.95 with a 0.05 frequency step. Each simulation represents a population of two drivers that are absolutely linked (*r =* 0, **C** and **D**) or unliked (*r =* 0.5, **E** and **F**) and have a low (*t =* 0.*2*, **C** and **E**) or high transmission bias (*t =* 1, **D** and **F**). The spread of two drivers is delayed by recombination as the gametes carrying one driver can be destroyed by the alternate driver. Strong drivers can go extinct in the presence of recombination, particularly when the starting frequency of the *wtfA*+ *wtfB*+ genotype is low (**E**; Table 2).

We also modeled the evolution of two drivers of differing strength both in the presence and absence of recombination (Figure 6A). Similar to our results with drivers of equal strength, we found that both drivers (i.e., the *wtfA*+ *wtfB*+ genotype) were fixed. Unlike the drivers of equal strength, however, there were no exceptional cases in which the *wtfA*+ *wtfB*+ genotype is not fixed when the two loci recombine (Table, 2; See Suppl. material for mathematical proof). The *wtfA*+ *wtfB*+ fixation rate was not dramatically affected by recombination rate (Figure 6A) and the stronger driver of the pair generally fixes faster (Figure 6B).

**Figure 6.**
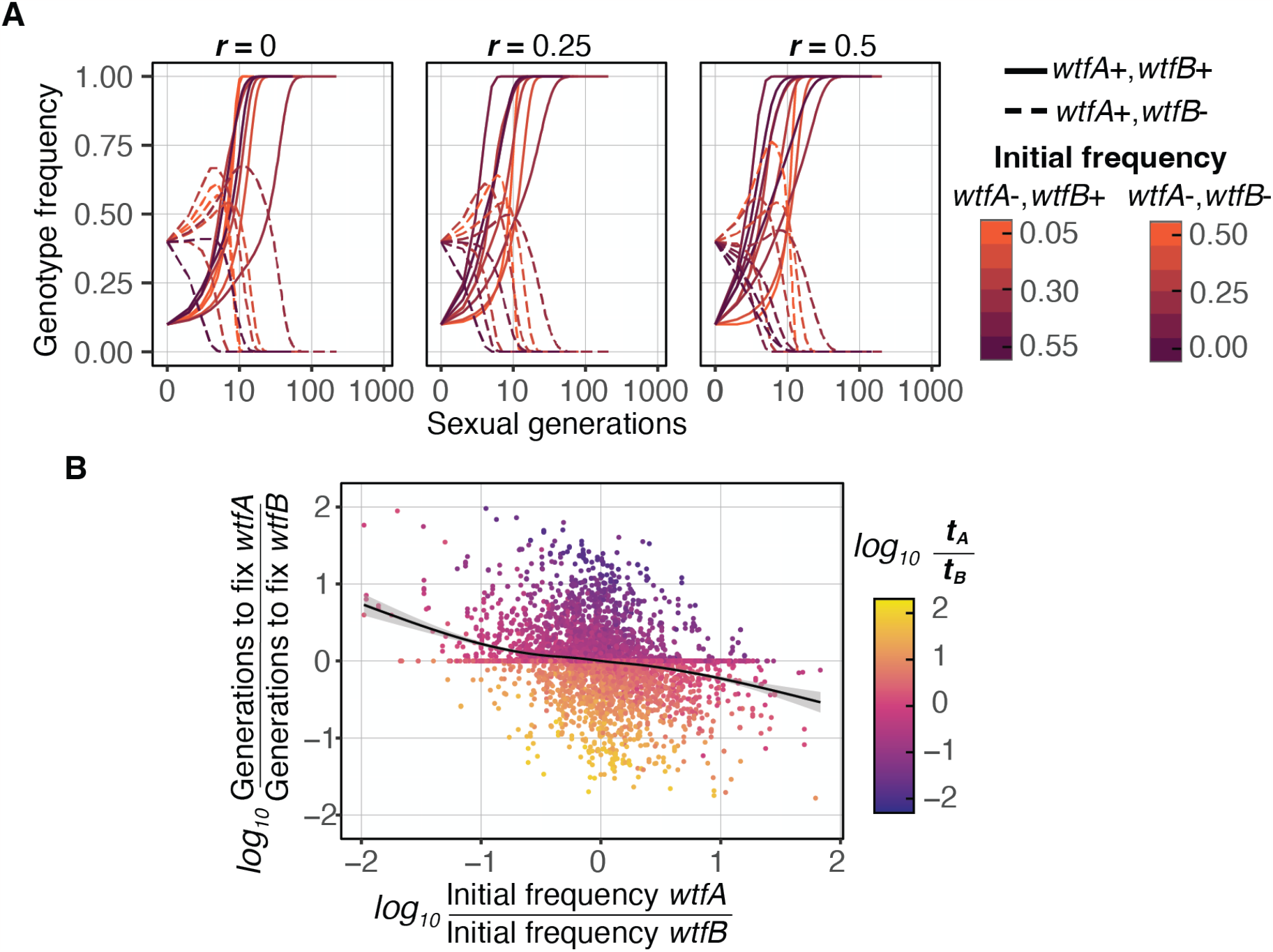
Drivers with larger transmission advantage tend to fix faster in a population. **A**) Simulations with varying transmission advantages *t*_*A*_ and *t*_*B*_ for absolutely linked (*r =* 0), mildly linked (*r =* 0.*2*5) or unlinked (*r =* 0.5) loci. The genotype frequencies of *wtfA+ wtfB*+ (solid, 0.1), *wtfA+ wtfB-* (dashed, 0.40) with varying *wtfA- wtfB-* initial frequencies with 0.1 steps. The remainder of each population is comprised of the *wtfA*- *wtfB*+. The genotype *wtfA+ wtfB+* goes to fixation when present (see Suppl. material for mathematical proof). The genotype of the double driver can decrease in the presence of recombination, but it eventually spreads to fixation. **B)** 10,000 initial populations were simulated with multiple recombination frequencies (*r =*0, 0.1, 0.2, 0.3, 0.4 and 0.5). The number of generations to fix a driver allele (i.e., *wtfA*) were compared to generations required to fix a second driver allele (i.e., *wtfB*). The stronger driver (larger *t*) tends to fix faster than a weaker driver, except in some cases when the weaker driver is more prevalent in a population. The black line is a local regression between X and Y axis. The shaded area is the standard error in the regression.

### Evolution of multiallelic drive loci

The *wtf* genes diverge so rapidly that different natural isolates of *S. pombe* can encode distinct drivers at a given locus (Eickbush et al. 2019). We therefore wanted to explore the evolution of multiple, distinct *wtf* drivers found at a single locus (eq. 3). When two driver alleles are considered, we found that the stronger driver generally spreads at the expense of the weaker driver, even if it is initially present at lower frequency. However, a weaker driver can drive a stronger driver to extinction if the starting frequency of the weaker driver is sufficiently high (Figure 7A; Table 2). Surprisingly, the steady states in which the population remains polymorphic is unstable (Figure 7A-B; Table 2).

**Figure 7.**
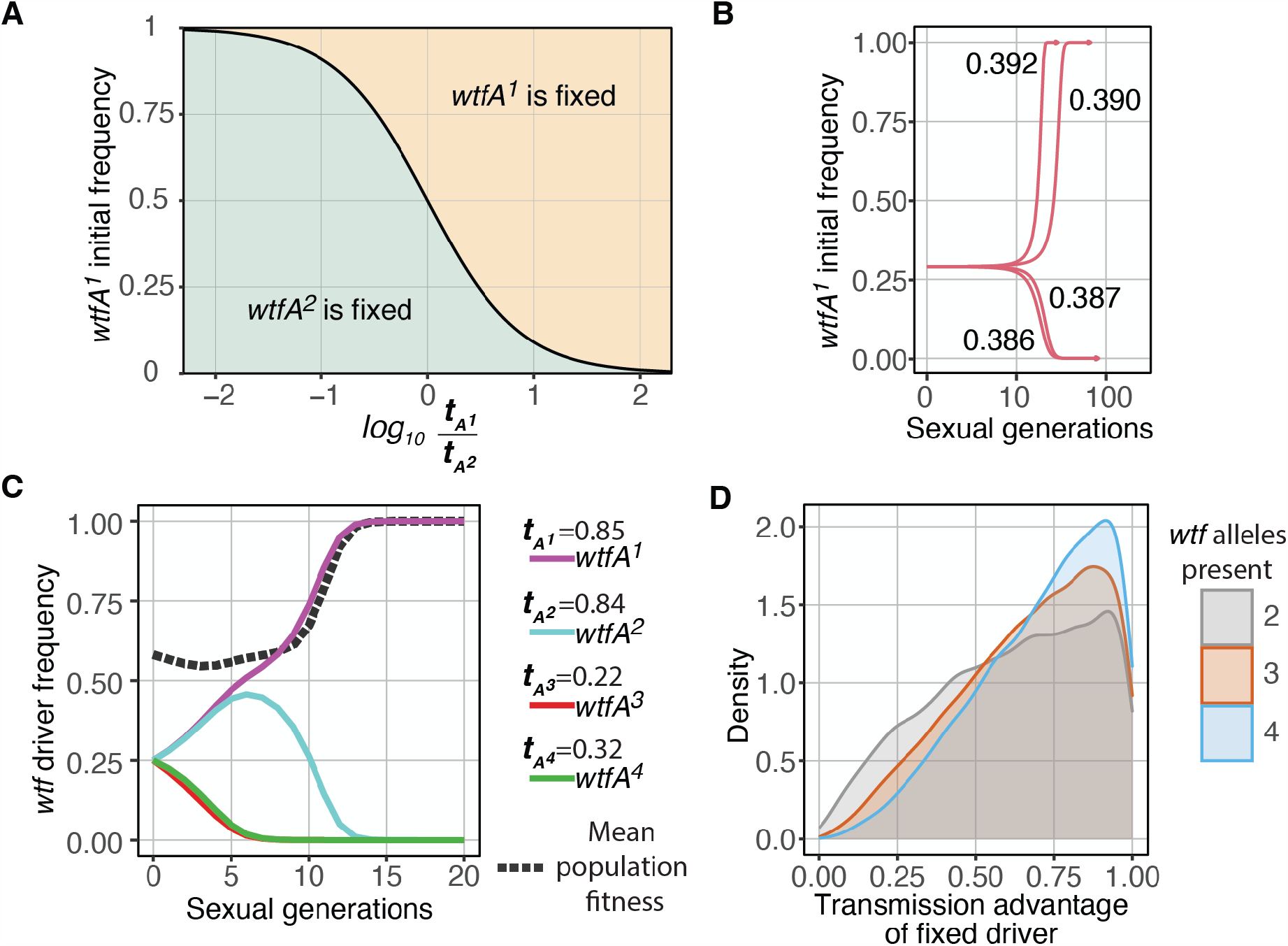
Evolution of populations with multiple allelic or absolutely linked *wtf* variants. **A)** Populations with only *wtfA*^*1*^ and *wtfA*^*2*^ drivers are considered to represent two alternate driving alleles of varying relative strengths. The plotted line (black) represents a steady state where the driver frequencies remain constant. At points above the line, the *wtfA*^*1*^ spreads to fixation. At points below the line, the *wtfA*^*2*^ *driver* spreads. The weaker driver can spread to fixation if the weaker driver starts in excess. **B)** The allele frequency changes over time for a few starting values near the limits illustrated in **A** (solid line). The starting frequency of the *wtfA*^*1*^ driver was 0.29 and the limit ratio of the drive strengths 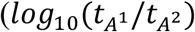 was 0.388. **C)** In populations with multiple allelic drivers or multiple absolutely linked driver on distinct haplotypes, the strongest driver (highest *t*) spreads to fixation when all alleles start at equal frequencies. The drive strength (*t*) of each driver is indicated on the key at the right. **D)** Depiction of 10,000 populations simulated with random initial frequencies for two, three, or four distinct allelic drivers (or absolutely linked drivers on distinct haplotypes). The transmission advantage of each driver was randomly selected. The histogram shows the distribution of transmission advantages for the fixed drivers for each set of simulations. The density axis indicates the proportion of events relative to 1. Drivers with high transmission advantages (t>0.9) are up to two times more likely to be fixed than drivers with an approximated medium transmission advantages 0.24, 0.43 and 0.55 for two, three, and four drivers competing, respectively.

When four drivers are present in the starting population at equal frequencies, strong drivers spread to then fix only the strongest driver (Figure 7C). In fact, the fixation of the strongest driver is the most common outcome under a range of drive strengths and starting allele frequencies as long as both drivers do not kill 100% of the gametes they inherit, 0 < *t* < 1 (Figure 7D).

## DISCUSSION

One route to accumulate drivers within a genome could be to fix them sequentially over time. If the drivers are independent, the evolutionary dynamics of this process would be no different than single driver evolution scenarios. However, in some species, drivers are polymorphic (Eickbush et al. 2019; Hall and Dawe 2018; Muirhead and Presgraves 2021; Vogan et al. 2019). To better understand the evolution of such duplicates, we modeled the evolution of duplicate killer meiotic drivers.

Our goal was to better understand the dynamics of meiotic driver duplicates in general. We used the *wtf* drivers of *S. pombe* as a model. This was a strength in that the parameters describing the behavior of *wtf* drivers in the lab are known and previous modeling matched well to laboratory experimental evolution analyses (Lopez Hernandez et al, 2021). Our study is, however, limited because we assumed an infinitely large, randomly mating population. These parameters do not describe all populations. For example, *S. pombe* grows clonally and cells are only passively mobile, both of which disfavor outcrossing. In addition, some isolates of *S. pombe* inbreed, even in the presence of potential outcrossing partners (Lopez Hernandez et al. 2021). We anticipate that inbreeding would slow, but not prevent, the fixation of two drivers (Lopez Hernandez et al, 2021). Drift, however, particularly combined with inbreeding, would likely significantly diminish the number of conditions under which the two drivers fix with high probability.

Our results have implications for understanding the evolution of natural drive systems, particularly poison-antidote killer meiotic drivers. Specifically, duplicates of such drive loci can be maintained or spread in a population under a broad range of conditions. This helps explain how the *wtf* genes have expanded in *Schizosaccharomyces* species. Similarly, isolates of *Podospora anserina* contain between zero and three distinct *Spok* drivers (Vogan et al. 2019). Like the *wtf* drivers, the *Spok* drivers are encoded in a single gene, which likely facilitates their establishment after being duplicated (Vogan et al, 2021). Partial duplication of poison-antidote drive systems in the form of antidote duplications have also been observed. For example, the first identified drive locus in the model plant *Arabidopsis thaliana*, contains multiple copies of the *APOK3* gene, which encodes an antidote to an unidentified poison (Simon et al. 2022). Although the impact of *APOK3* duplications is unknown, such antidote duplications could potentially make a driver more efficient by ensuring extra protection for meiotic products that inherit the drive locus.

The duplication of drivers that do not use a poison-antidote mechanism may be relatively more constrained. For example, chromosome ‘knobs’ in maize drive by preferential segregation into the egg cell during female meiosis (Dawe et al. 2018; Novitski 1957). Drive of knobs is affected by chromosomal position, which likely constrains the evolution of duplicated knob sequences (Swentowsky et al. 2020). Knobs are also quite large, which may also limit their duplication potential. Despite these factors, multiple knobs are found on most maize chromosomes (Hufford et al. 2021).

Similarly, killer-target drive systems are also likely more constrained in their duplication. These drivers use a killer element to destroy the meiotic products that inherit a target locus that is found on the competing haplotype but is not found on the driving haplotype. Duplications of a killer to a location not absolutely linked in cis to the parent locus would likely be lost as the duplicate would not benefit from drive and would sometimes be destroyed by drive. However, duplications of the killer element linked in cis to the original drive locus could be favored if duplications strengthened the drive of the haplotype (Crow 1991). For example, an X chromosome-linked killer that targeted gametes inheriting the Y chromosome could duplicate on the X chromosome to enhance drive of the X. Although the mechanisms of drive are not yet known, X-linked expansions of drive genes have been observed (Kruger et al. 2019; Muirhead and Presgraves 2021; Vedanayagam et al. 2021).

Finally, this work has implications that could be considered in the design of synthetic gene drives to spread desirable traits in a population (Burt and Crisanti 2018). Single-gene poison-antidote meiotic drivers, like the *wtf* drivers, are an attractive candidate component for such synthetic gene drives. Their strong drive, small size, autonomy, and inability for the critical drive components to be uncoupled by recombination are all ideal for promoting the spread of a desired locus or chromosome in a population. Unfortunately, those same features also increase the possibility that a gene drive could spread within a genome. Such duplication could lead to less predictable control and other undesirable outcomes. As discussed above, killer-target meiotic driver systems have less duplication potential and thus may be better guides for engineering gene drives to spread desirable traits in a population, but not within genomes.

