## Supplementary material for "Modeling the Evolution of Populations with Multiple Killer Meiotic Drivers"

The purpose of this section is to explore the mathematical stability of the steady states (e.g. fixation of a genotype) described in the main text using eigenvalues of the Jacobian matrix. In lay terms, we used the math described below to determine if the population will return to the steady state position upon perturbation. The conclusions we draw from the analyses are summarized in Table 2. The first line of Table 2 considers a general case where there are no restrictions on the values of $t$ or $r$ and *wtfA* and *wtfB* are distinct drivers (as in Figures 4 and 5). Under those conditions, there are four possible steady state solutions to consider corresponding to fixation of the four different genotypes ($x_1$-$x_4$). Fixation of the *wtfA*+ *wtfB*+ genotype ($x_1$=1) stable because after perturbing that population by adding other genotypes, the population would evolve until *wtfA*+ *wtfB*+ was fixed again. Fixation of all other genotypes (i.e. $x_2$-$x_4$) is not stable.

The second line of Table 2 describes the unstable conditions that are illustrated in Figure 5F where either the *wtfA*+ *wtfB*+ ($x_1$) or the *wtfA*- *wtfB*- ($x_4$) genotype is fixed. The third line of the table describes the unstable conditions illustrated in Figure 7A-7B where either one of two genotypes is fixed, depending on the parameters considered.

The eigenvalues of the Jacobian matrix for all four recurrence equations (eq. 1.1,2.2,2.3,2.4) determine the stability of solutions in a system of equations. A solution is stable only when the (real part of) leading eigenvalue is less than 1 and unstable when it is greater than 1. In cases where the associated eigenvalues are exactly one, the solution stability cannot be defined by the Jacobian matrix alone.

In the solutions that maintained a polymorphic population, either by maintaining a double driver genotype *wtfA*+ *wtfB*+ and a non-driver genotype *wtfA*- *wtfB*- or by keeping only the two single driver carriers *wtfA*+ *wtfB*- and *wtfA*- *wtfB*+ genotypes in populations were unstable (Table 2), determined by the Jacobian. The other solutions only included the fixation of one genotype. If genotypes that do not carry both drivers $x_2$, $x_3$, $x_4$ are perturbed by the presence of the double driver genotype, it leads to the fixation of the *wtfA*+ *wtfB*+ genotype, $x_1 = 1$ . The solution for the fixation of the $x_1$, however, cannot be fully determined by the Jacobian matrix. The leading eigenvalue of this solution is 1. This uncertainty is addressed in the following text.

To fully determine the stability of this solution, we considered a system of N discrete time dynamical equations,

$$x_i[n+1] = f_i(x_1[n], x_2[n], \ldots, x_N[n])$$

where $1 \leq i \leq N$ is the number of the variable $x_i$, corresponding to each "$i$" genotype, and $n =$ 0,1,2, .... Is the time step number (generation $n$).

We use the vector notation for the functions $F = \{f_1, f_2, \ldots, f_N\}$ and the variables $X[n] = \{x_1[n], x_2[n], \ldots, x_N[n]\}$ and rewrite the equation as, $X[n+1] = F(X[n])$. This system of equations calculates the N genotypes to the next generation as a function of the previous generation. The steady state (SS) solutions $\tilde{x}_i$, satisfy the equations $\tilde{x}_i = f_i(\tilde{x}_1, \tilde{x}_2, \ldots, \tilde{x}_N)$ also represented as $\tilde{x} = F(\tilde{x})$ for the system. In the case where the double driver genotype is fixed $\tilde{x}_1 = \{1,0,0,0\}$.

To determine the stability of that solution we perturbed the frequency of the rest of genotypes, at the expense of fixed double driver genotype, with a deviation $\delta$. Such deviation then satisfies that the change in the fixed driver, $\delta_1$, is equal to the sum of in the increase of all the other genotypes $-(\delta_2 + \delta_3 + \delta_4)$. The perturbation of the solutions is thus $\delta_1 = -(\delta_2 + \delta_3 + \delta_4) = -\delta$.

By using a small deviation $\delta_i[n] \ll 1$ from the SS, we write that the solution $x_i[n] = \tilde{x}_i + \delta_i[n]$, and substitute it into the original equations. We have the solution of the system $\tilde{x} + \delta X[n+1] = F(\tilde{x} + \delta X[n])$. By expanding the right hand side of the equation into the Taylor series up to the linear terms and considering that in Jacobian matrix elements $J_{ij} = \partial f_i / \partial x_j$ are defined as partial derivatives of the functions $f_i$ with respect to variables $x_j$ we find $\delta X[n+1] = J\delta X[n]$. In this notation it is possible to observe that the dynamics of the deviation vector $\delta X$ is determined completely by the properties of the Jacobian matrix, namely, its eigenvalues $\lambda_i$ and the eigenvectors $e_i$, associated with each solution.

The deviation from the steady state, $\delta X[n]$, can be expressed as an expansion $\delta X[n] = \sum_{i=1}^{N} r_i[n] e_i$ where $r_i$ is the contribution of each eigenvector $e_i$. This relation implies that $\delta X[n+1] = \sum_{i=1}^{N} r_i[n+1] e_i$. On the other hand, we observe that $J\delta X[n] = \sum_{i=1}^{N} r_i[n] J e_i = \sum_{i=1}^{N} r_i[n] \lambda_i e_i$. Now using the equality $\delta X[n+1] = J\delta X[n]$ we obtain a relation $r_i[n+1] = \lambda_i r_i[n]$, where the coefficient dynamics is completely determined by the eigenvalue associated with each solution.

The eigenvalues $\lambda_i$ show how the expansion coefficients $r_i$ change with every time step. If $|\lambda_i|>1$ the coefficient $r_i$ increases and thus the deviation in the direction set by $e_i$ grows. If $|\lambda_i|<1$ the coefficient $r_i$ decreases and thus the deviation in the direction $e_i$ shrinks. Finally, if $|\lambda_i|=1$ the coefficient $r_i$ stays constant and thus the deviation in the direction $e_i$ does not change.

In our case for the solution $\tilde{x}= \{1,0,0,0\}$ we have $\lambda_1=1$ and $\lambda_2$, $\lambda_3$, $\lambda_4<1$. After sufficiently large number of time steps we will have $r_2 = r_3 = r_4 \rightarrow 0$, while $r_1$ remains constant with $e_1 = \{1,0,0,0\}$. After introducing the initial perturbation in the system $\delta X[0] = \{\delta_1, \delta_2, \delta_3, \delta_4\}$ and since $\delta_1 = -\delta = -(\delta_2 + \delta_3 + \delta_4)$, now we can track the dynamics of the deviation $\delta X[n]$ starting from its initial value $\delta X[0] = \{-\delta, \delta_2, \delta_3, \delta_4\}$. Any variation of the solution where the double driver genotype is fixed will show how much it deviates or approached the original solution $\tilde{x}_1 = \{1,0,0,0\}$.

We use the equation $\delta X[0] = \sum_{i=1}^{N} r_i[0] e_i$ with N=4 to find the value $r_1 = r_1[0]$. The direct computation shows that $r_1[0] = -\delta$ . This allows to write for the deviation $\delta X_1 \rightarrow r_1 e_1 = \{-\delta, 0,0,0\}$ leading to $X \rightarrow \{1-\delta, 0,0,0\}$. Considering that $x_1 + x_2 + x_3 + x_4 = 1$ at all times we conclude that the perturbation $\delta \rightarrow 0$, thus the perturbed frequencies for each genotype, as steps increase, approach the only stable solution $\tilde{x}_1 = \{1,0,0,0\}$. The double driver genotype will get fixed even in the presence of any other genotype. This solution stability is not affected by elimination of any variable $x_i$ reducing by one the number of equations in the system.
